# Analysis of LRRC45 indicates cooperative functions of distal appendages at early steps of ciliogenesis

**DOI:** 10.1101/205625

**Authors:** Bahtiyar Kurtulmus, Cheng Yuan, Jakob Schuy, Annett Neuner, Shoji Hata, Georgios Kalamakis, Ana Martin-Villalba, G. Pereira

## Abstract

Cilia perform essential signalling functions during development and tissue homeostasis. Ciliary malfunction causes a variety of diseases, named ciliopathies. The key role that the mother centriole plays in cilia formation can be attributed to appendage proteins that associate exclusively with the mother centriole. The distal appendages form a platform that docks early ciliary vesicles and removes CP110/Cep97 inhibitory complexes from the mother centriole. Here, we analysed the role played by LRRC45 in appendage formation and ciliogenesis. We show that the core appendage proteins Cep83 and SCLT1 recruit LRRC45 to the mother centriole. Once there LRRC45 recruits FBF1. The association of LRRC45 with the basal body of primary and motile cilia in differentiated and stem cells reveals a broad function in ciliogenesis. In contrast to the appendage components Cep164 and Cep123, LRRC45 was neither essential for docking of early ciliary vesicles nor for removal of CP110. Rather, LRRC45 promotes cilia biogenesis in CP110-uncapped centrioles by organising centriolar satellites and promoting the docking of Rab8 GTPase-positive vesicles. We propose that, instead of acting solely as a platform to recruit early vesicles, centriole appendages form discrete scaffolds of cooperating proteins that execute specific functions that promote the initial steps of ciliogenesis.

## Introduction

The primary cilium is a microtubule-based organelle that protrudes from the surface of several cell types in the human body. It has mechanosensory and signaling functions due to its association with receptors of important signaling pathways, such as hedgehog (Shh) and Wnt (Goetz and Anderson, 2010). Defects in primary cilia formation or function are the underlying cause of severe genetic diseases, collectively named ciliopathies, in which embryonic development and/or functioning of multiple organs in the human body can be affected (Reiter and Leroux, 2017).

The primary cilium emerges from the mother centriole that converges into the basal body of the cilium. Each centrosome is composed of two centrioles, cylindrical structures consisting of a nine-fold array of triplet microtubules, and pericentriolar material (PCM). The two centrioles of a centrosome are different in function and composition. The mother but not daughter centriole associates with an array of proteins, named appendage proteins, that upon cell cycle exit gives rise to the cilium (Nigg and Stearns, 2011).

Ciliogenesis is a multi-step process that requires the concerted action of several components, including mother centriole-associated proteins and the vesicular transport machinery, which together are important for the establishment and maintenance of the ciliary compartment (Sanchez and Dynlacht, 2016). At early stages of ciliogenesis, the mother centriole associates with small ciliary vesicles, most likely derived from the Golgi-apparatus (Das and Guo, 2011; Qin, 2012). After docking to the mother centriole, small vesicles are fused thereby forming a large ciliary vesicle that caps this centriole (Sanchez and Dynlacht, 2016). The fusion process depends on the membrane-shaping proteins (EHD1, EHD3) as well as the SNARE component SNAP-29 (Lu et al., 2015). Ciliary membrane establishment also requires components of the Rab-GTPase cascade, which includes the GTPase Rab11, the small Rab GTPase Rab8 and the Rab8 activator Rabin8 (Knodler et al., 2010; Nachury et al., 2007; Yoshimura et al., 2007). Rabin8 and Rab8 accumulate at the centrosomes shortly after induction of ciliogenesis implying a local activation of Rab8 to promote ciliary vesicle formation (Westlake et al., 2011). Alongside the process of ciliary vesicle establishment, the CP110/Cep97 complex is removed from the mother centriole by targeted protein degradation (Spektor et al., 2007). CP110/Cep97 removal allows the extension of microtubules to form the ciliary axoneme (Kobayashi et al., 2011). The intraflagellar transport (IFT) machinery also controls cilia formation and extension (Follit et al., 2006; Jurczyk et al., 2004; Pazour et al., 2002; Pazour et al., 2000; Pedersen and Rosenbaum, 2008). In addition, large cytoplasmic protein complexes, so called centriolar satellies, are involved in the transport of building blocks to the basal body (Craige et al., 2010; Garcia-Gonzalo et al., 2011; Hori et al., 2014; Jin et al., 2010; Klinger et al., 2013; Kurtulmus et al., 2016).

Distal appendage proteins at the mother centriole are key for initial steps of cilia biogenesis. Distal appendage components include the C2-calcium-dependent domain protein C2CD3, Cep83/CCDC41, the sodium channel and clathrin linker protein 1 (SCLT1), Cep123/Cep89, Fas Binding Factor 1 (FBF1) and Cep164 (Graser et al., 2007; Joo et al., 2013; Schmidt et al., 2012; Sillibourne et al., 2013; Tanos et al., 2013; Wei et al., 2013; Ye et al., 2014). The protein C2CD3 positively controls centriole length (Thauvin-Robinet et al., 2014). In the absence of C2CD3, centrioles lack Cep83 and, consequently, all additional distal components, given that Cep83 is necessary for binding of Cep123, SCLT1, Cep164 and FBF1 to the mother centriole (Tanos et al., 2013; Ye et al., 2014). Upon depletion of distal components, including Cep164, Cep123 and Cep83, the initial docking of small ciliary vesicles at the mother centriole does not occur. In addition, the inhibitory CP110/Cep97 complex is not displaced from the mother centriole and axoneme extension is blocked. Distal appendages interact with components of the vesicular transport machinery, including Rab8, as well as with protein kinases and phosphatases that influence CP110/Cep97 recruitment (Bielas et al., 2009; Cajanek and Nigg, 2014; Goetz et al., 2012; Humbert et al., 2012; Kuhns et al., 2013; Oda et al., 2014; Schmidt et al., 2012; Tanos et al., 2013; Xu et al., 2016; Ye et al., 2014). Therefore, the current model is that distal appendages are indispensable for vesicle docking at the mother centriole and all subsequent steps thereafter.

Despite the importance of appendage proteins for cilia biogenesis, it is not fully understood how appendages are formed and how they regulate initial steps of ciliogenesis. Here, we investigated the function of the leucine-rich repeat protein, LRRC45, in cilia formation. LRRC45 was reported to be required for centrosome cohesion at the proximal end of centrioles, as part of the linker that holds duplicated centrosomes together (He et al., 2013). LRRC45 is recruited to the proximal end of the centrioles by the protein C-Nap1 (He et al., 2013). In addition, a pool of LRRC45 was shown to associate with mother centriole appendages (He et al., 2013), implying a direct function in ciliation. We now show that LRRC45 associates with the distal appendages of the mother centriole in a Cep83 and SCLT1-dependent manner. Depletion of LRRC45 impaired ciliogenesis independently of C-Nap1. We show that LRRC45 is required for Rab8 and satellite recruitment to centrosomes but not for the CP110/Cep97 complex removal or small ciliary vesicle docking at the mother centriole. We propose that LRRC45 constitutes a novel branch of distal appendages that works in parallel to Cep164 and Cep123 to control distinct steps at early stages of ciliogenesis.

## Results

### LRRC45 contributes to cilia formation independently of C-Nap1

LRRC45 was shown to associate with the basal body of primary cilia in retina pigment epithelial (RPE1) cells (Figure 1A) (He et al., 2013). We also observed that LRRC45 associates with the basal body of ciliated murine fibroblasts, neural stem cells and motile cilia of ependymal cells (Figure 1A-B). This indicates that LRRC45 may have a function in promoting ciliogenesis in several cell types, including cells with motile cilia.

**Figure 1.**
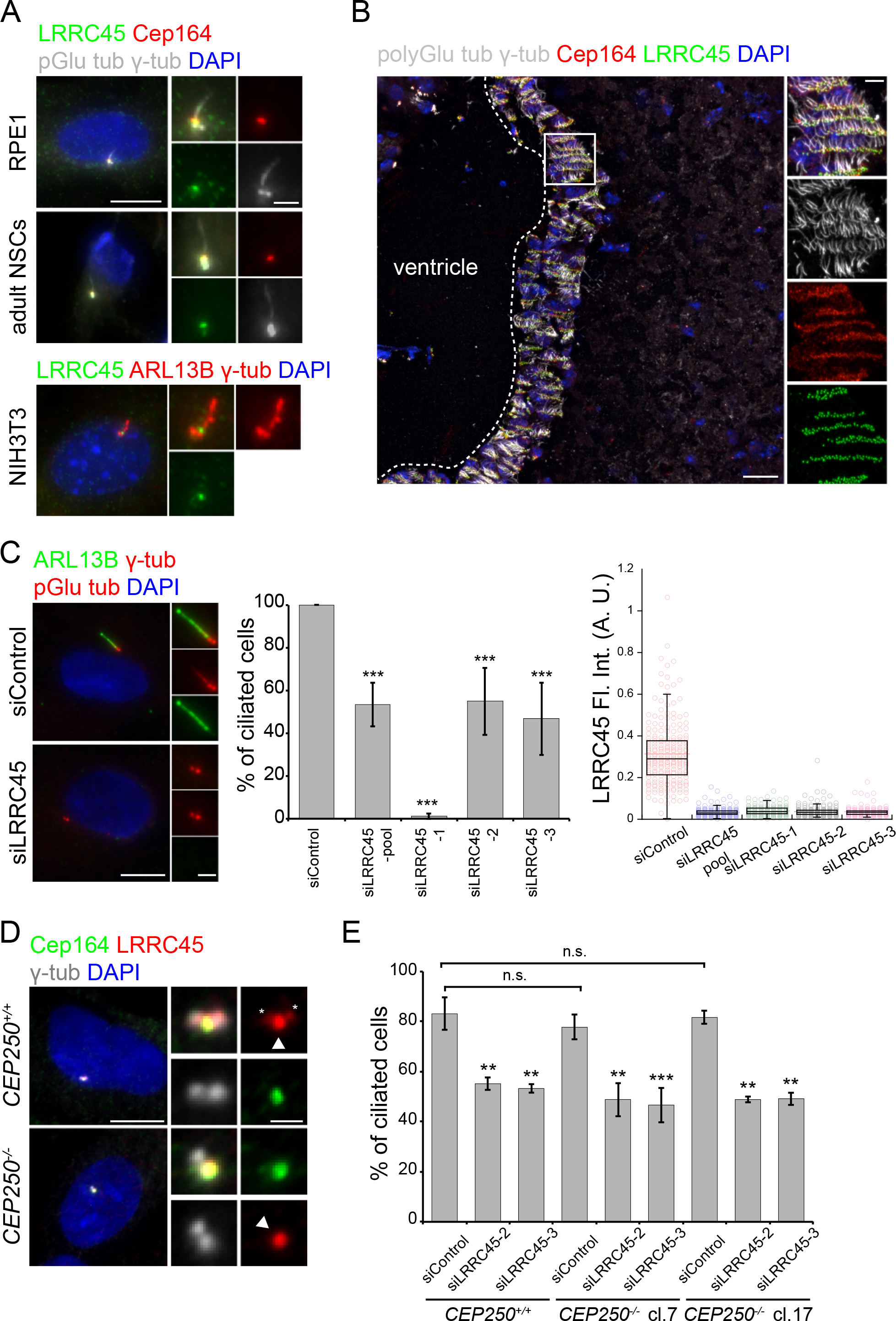
LRRC45 localizes to the basal body of ciliated cells and is required for cilia formation. (A) Representative images showing localisation of LRRC45 (green) with distal appendage protein Cep164 (red) at the basal body in ciliated RPE1 cells and adult NSCs, poly-glutamylated (pGlu) and γ-tubulin served as markers for cilium and basal body, respectively (upper panel). LRRC45 localisation at the basal body in ciliated NIH-3T3 cells. ARL13B and γ-tubulin served as markers for cilium and basal body, respectively (lower panel). Scale bars: 10 μm and 2 μm. (B) LRRC45 (green) co-localisation with Cep164 (red) in multi-ciliated ependymal cells lining the ventricle of adult mouse brain. Poly-glutamylated (pGlu) and γ-tubulin served as markers for cilium and basal body respectively. Blue colour represents nuclear staining by DAPI. Magnifications of the centrosome area or indicated region (in B) are shown to the right. Scale bars: 20 μm and 5 μm. (C) Representative images and quantification of ciliated and non-ciliated cells upon treatment of RPE1 cells with control- or indicated LRRC45-siRNAs and serum starved for 48 h. ARL13B (green) served as a ciliary membrane marker, poly-glutamylated (pGlu tub) and γ-tubulin served as markers for ciliary axoneme and basal body, respectively. DAPI stained nuclei. Scale bars: 10 μm and 2 μm. The bar graph indicates the mean ± standard deviation of three independent experiments, n ≥ 100 for each condition in each experiment. The box/dot plots show LRRC45 fluorescent intensity (arbitrary unit) measured at the centrosome in control and LRRC45 depleted cells. The graph is a representative of one experiment out of three. Number of centrosomes quantified in this particular experiment are 218, 231, 183, 180 and 176 for siControl, siLRRC45 pool, siLRRC45-1, -2, and -3, respectively. *** p < 0.001 (Student’s t-test). (D) Representative images showing LRRC45 (red) localisation in WT and *CEP250*^-/-^ RPE1 cells. Cep164 (green) γ-tubulin (gray) served as markers for mother centriole and centrosomes, respectively. DAPI stained nuclei. The arrowhead marks LRRC45 that co-localises with Cep164, while the two asterisks mark LRRC45 at the proximal ends of centrioles in *CEP250*^+/+^ cells. Scale bars: 10 μm and 1 μm. (E) Bar graph shows percentage of ciliated cells after 48 h of serum starvation in *CEP250*^+/+^ and *CEP250*^-/-^ (clones 7 and 17) RPE1 cells upon treatment with control- and the indicated LRRC45 siRNAs. Results show the mean ± standard deviation of three independent experiments, n ≥ 100 for each condition in each experiment. ** p<0.01 and *** p < 0.001 (Student’s t-test).

To understand the function of LRRC45 in ciliogenesis, we depleted LRRC45 using three independent small interfering RNAs (siRNA) and a pool of 4 siRNAs. Loss of LRRC45 led to reduction in cilia formation in RPE1 cells (Figure 1C). This effect was specific for LRRC45, as it could be rescued by the expression of murine *LRRC45* (Figure S1A). However, we were unable to rescue the strong cilia loss phenotype obtained with the published *LRRC45* siRNA sequence (LRRC45-1, Figure S1B) (He et al., 2013). This suggests that the LRRC45-1 siRNA has an off-target effect that exacerbates the cilia loss phenotype.

LRRC45 associates with the proximal end of the mother and daughter centrioles in dependence of the protein C-Nap1 (encoded by *CEP250*) (He et al., 2013). In addition, a fraction of LRRC45 associated with the mother centriole independently of C-Nap1 (Figure 1D). Because of this dual localisation, we asked which fraction of LRRC45 contributes to cilia formation. For this, we used *CEP250* KO cells (Panic et al., 2015). These cells can form cilia similarly to RPE1 wild type cells (Figure 1E). Depletion of LRRC45 in RPE1 *CEP250* KO cells caused the same degree of cilia loss phenotype as observed in RPE1 wild type cells (Figure 1E). We concluded that LRRC45 regulates ciliogenesis independently of its function at the centrosomal linker.

Recently, the daughter centriole has been implied in ciliogenesis due to its proximity to the mother centriole (Loukil et al., 2017). We therefore asked whether lack of cilia upon LRRC45 depletion in WT and *CEP250* KO RPE1 cells correlated with the distance between both centrioles (Figure 2). Under serum starvation, mother and daughter centrioles were significantly more separated in *CEP250* KO in comparison to wild type RPE1 cells (Figure 2A). Yet, *CEP250* KO cells were able to form cilia even when mother and daughter centrioles were more than 10.5 μm apart (Figure 2A). This indicated that the physical proximity of daughter and mother centriole is not required for ciliogenesis. Neither in cycling nor in serum starved RPE1 cells lacking LRRC45, we could observe an increase in the distance between mother and daughter centrioles irrespectively of whether these cells had a cilium or not (Figure 2B). Together, these data indicate that defective ciliation upon LRRC45 knockdown is most likely unrelated to mother-daughter centriole proximity. It also indicates that, in contrast to U2OS or HeLa cells (He et al., 2013), LRRC45 does not play a major role in centrosome cohesion in RPE1 cells. This might be explained by an alternative microtubule-dependent centrosome-cohesion pathway described to be present in RPE1 but to a lesser extent in cancer cell lines (Panic et al., 2015).

**Figure 2.**
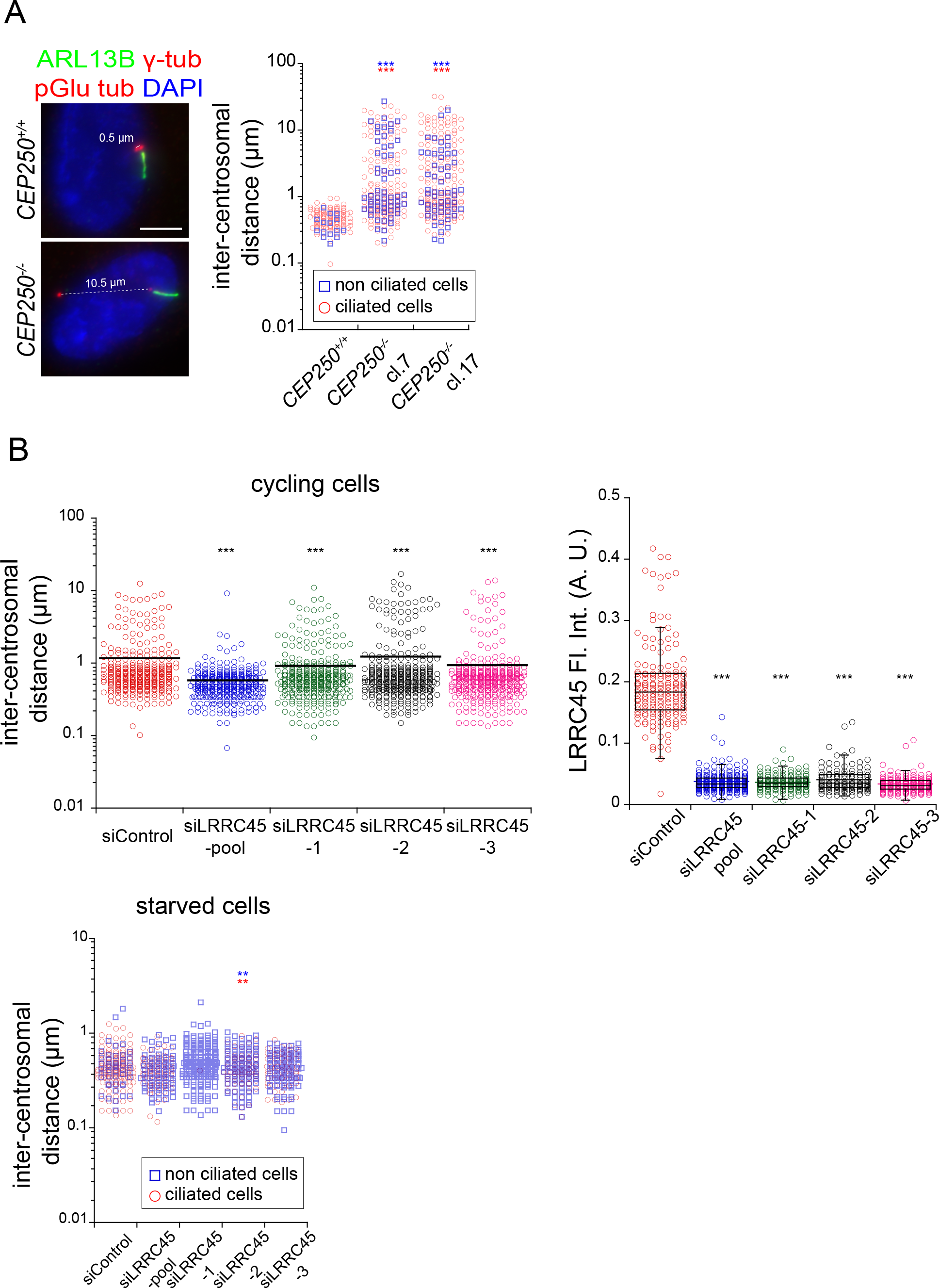
Depletion of LRRC45 does not cause premature centrosome separation. (A) Representative images showing 0.5 μm and 10 μm inter-centrosomal distance in ciliated WT and *CEP250*^-/-^ RPE1 cells, respectively. ARL13B (green), poly-glutamylated (pGlu tub) and γ-tubulin (red) served as markers for ciliary membrane, axoneme and basal body, respectively. The dot plots show the distribution of inter-centrosomal distances in ciliated (red circle) and non-ciliated (blue square) in WT and two independent clones of *CEP250*^-/-^ RPE1 cells after 48 h of serum starvation. Results from one out of three independent experiments are shown. Number of ciliated cells analysed are 125, 165, and 175 in WT, *CEP250*^-/-^ clone 7 and clone 17, respectively. Number of non-ciliated cells analysed are 16, 58, and 55 in WT, *CEP250*^-/-^ clone 7 and clone 17, respectively. *** p<0.001 (unpaired Wilcoxon-Mann-Whitney Rank Sum Test). Red and blue asterisks represent the significant increase in the inter-centrosomal distances between WT and *CEP250*^-/-^ clones. (B) Dot plots showing inter-centrosomal distances upon LRRC45 depletion in cycling (left) and serum starved (right) cells. Plots of cycling cells are a combination of three independent experiments with total n of 315, 332, 348, 353, and 353 for each respective siRNA. The box/dot plots show LRRC45 fluorescent intensity (arbitrary unit) measured at the centrosome in control and LRRC45 depleted cells. The graph is a representative of one experiment out of three. Number of centrosomes quantified in this particular experiment are 164, 216, 137, 111 and 155 for siControl, siLRRC45 pool, siLRRC45-1, -2, and -3, respectively. Plots of starved cells show the distribution of inter-centrosomal distances in ciliated (red circles) and non-ciliated (blue squares) upon LRRC45 knock down. It is a combination of two independent experiments with total n of 149, 73, 1, 70, 63 for ciliated cells and 46, 99, 198, 124, 164 for non-ciliated cells for each respective siRNA (from left to right). ** p<0.01 and *** p < 0.001 (unpaired Wilcoxon-Mann-Whitney Rank Sum Test). Red and blue asterisks represent the significant increase of inter-centrosomal distance between control and LRRC45 depletion.

### Cep83 and SCLT1 direct LRRC45 to distal appendages

Our data indicates that the LRRC45 pool at centriole appendages contributes to ciliogenesis (Figure 1-2). Using specific LRRC45 antibodies (Figure S2), we determined LRRC45 sub-cellular localisation by super-resolution microscopy (Figure 3). Three dimensional-structured illumination microscopy (3D-SIM) resolved LRRC45 localization in top view images of the mother centriole as a ring-like structure with a diameter of 264+/-31 nm (Figure 3A). The size of the ring formed by LRRC45 was significantly smaller than the one formed by the distal appendage protein Cep164 (350+/-36 nm) but similar to the rings of the distal component Cep123 (288+/-49 nm) or the sub-distal appendage protein ODF2 (260+/-32 nm) (Figure 3A). Analysis of pairwise co-localisation between LRRC45 and distal (Cep123, SCLT1, Cep83 and Cep164) or sub-distal (ODF2) components by stimulated emission depletion (STED) microscopy revealed that LRRC45 appeared as dotted ring-like structures around centrioles that did not co-localise with ODF2 (Figure 3B). LRRC45 co-localised with Cep83 (Figure 3C) but it only partly colocalised with Cep164 or SCLT1 (Figure 3D). Interestingly, Cep123 could be resolved in some cells as one or two rings (Figure 3F, lower panel). In cells with two rings, LRRC45 localised between the two rings (Figure 3F), whereas in cells with one Cep123 ring, LRRC45 located mostly proximal to it (i.e. between Cep123 and sub-distal appendages).

**Figure 3.**
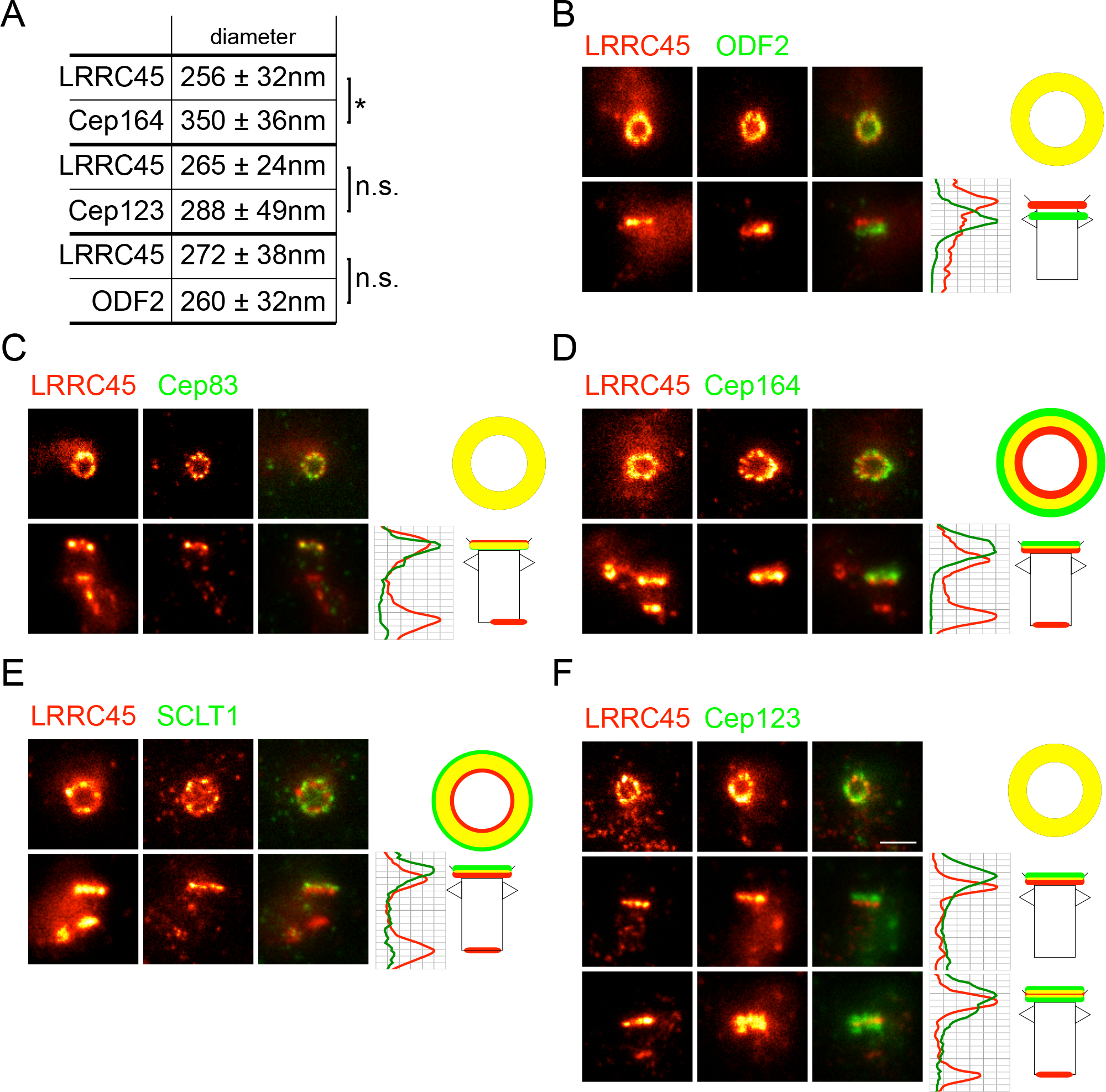
LRRC45 localizes to the distal but not sub-distal appendages. (A) Table showing mean diameter ± standard deviation of LRRC45 ring with respect to Cep164, Cep123 and ODF2. n are 13 (LRRC45 - Cep164), 7 (LRRC45 - Cep123) and 8 (LRRC45 - ODF2). Representative images of LRRC45 co-stainings with ODF2 (B), Cep83 (C), Cep164 (D), SCLT1 (E), Cep123 (F) from top view (top panels) and side view (bottom panels) using STED microscopy. Line graphs show the plot profile of LRRC45 (red) and respective appendage protein (green) from side view pictures. Sketches of top and side views represent co-localisation (yellow) of LRRC45 (red) and the indicated appendage protein (green). Scale bar: 0.5 μm.

Previously a hierarchical network of distal appendage proteins was unravelled by siRNA depletion experiments (Figure 4A). To position LRRC45 within this network, we first investigated its localisation upon depletion of appendage components. Cep83 targets the appendage proteins Cep123, Cep164, FBF1 and SCLT1 to the mother centriole (Tanos et al., 2013) (Figure S3). Centriole association of LRRC45 was significantly decreased upon depletion of Cep83 (Figure 4B) or SCLT1 (Figure 4C). In contrast, depletion of Cep123, Cep164 or FBF1 did not affect the centrosomal levels of LRRC45 (Figure S4A-C). LRRC45 localisation was also not influenced by depletion of ODF2 and vice-versa (Figure S4D and S6A). In conclusion, LRRC45 is mainly targeted to distal appendages via Cep83 and SCLT1.

**Figure 4.**
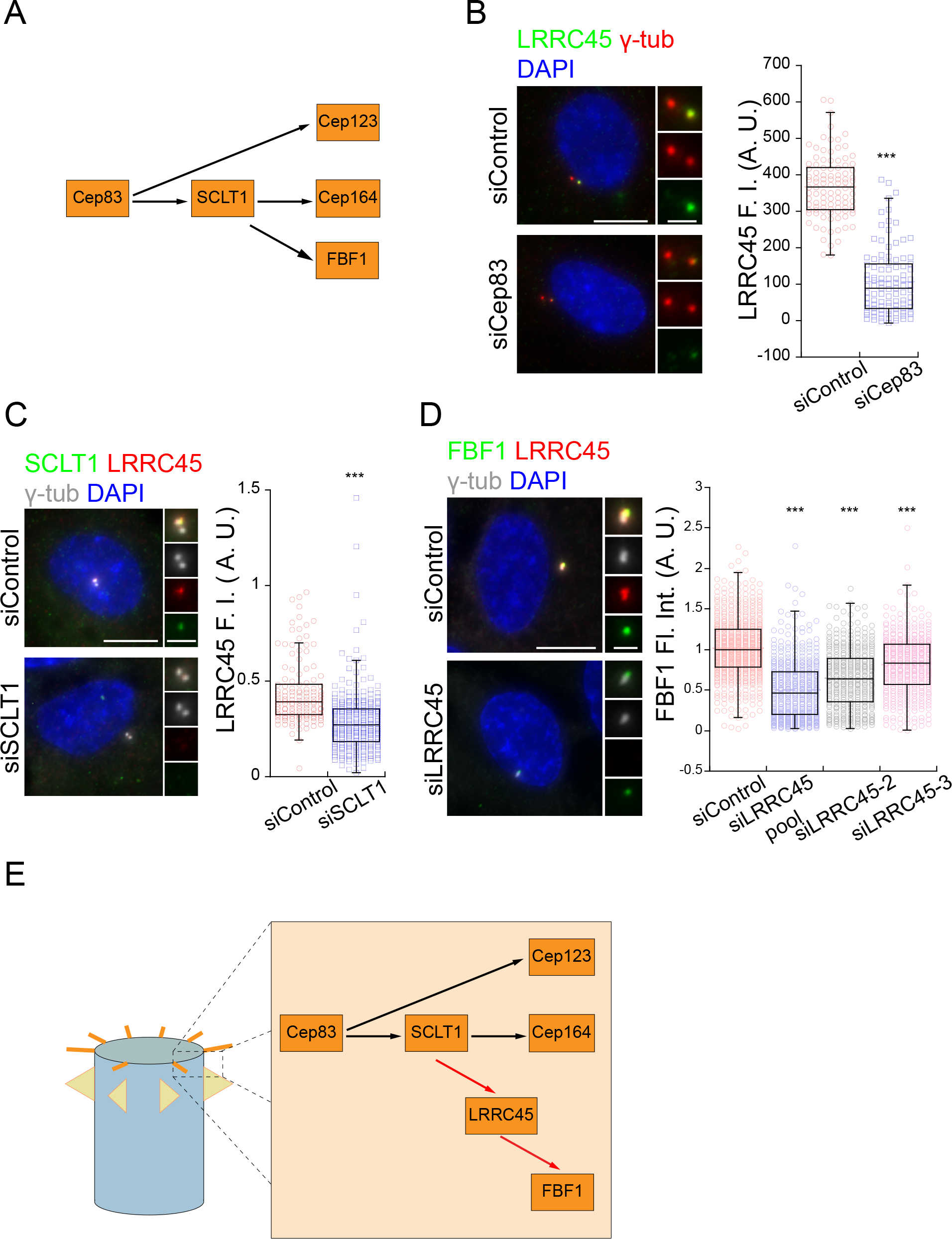
LRRC45 bridges Cep83-SCLT1 and FBF1 in the hierarchy of appendages. (A) Hierarchy of distal appendages as previously published (Tanos et al., 2013). (B) Representative images showing LRRC45 (green) localisation in control and Cep83 depleted RPE1 cells. γ-tubulin (red) and DAPI (blue) serve as markers for centrosomes and nuclei, respectively. Box/dot plot shows quantification of fluorescent intensities of LRRC45 on the centrosomes in control and Cep83 depleted cells (n, 101 for each condition). *** p < 0.001 (unpaired Wilcoxon-Mann-Whitney Rank Sum Test). (C) Representative images showing LRRC45 (red) and SCLT1 (green) localisation in control and SCLT1 depleted RPE1 cells. γ-tubulin (grey) and DAPI (blue) serve as markers for centrosomes and nuclei, respectively. Box/dot plots show quantification of fluorescent intensities of LRRC45 on the centrosomes in control and SCLT1 depleted cells (n: 126 and 266 for control and SCLT1 depletion, respectively). *** p < 0.001 (unpaired Wilcoxon-Mann-Whitney Rank Sum Test). (D) Representative images showing LRRC45 (red) and FBF1 (green) localisation in control and FBF1 depleted RPE1 cells. γ-tubulin (grey) and DAPI (blue) serve as markers for centrosomes and nuclei, respectively. Box/dot plots show quantification of fluorescent intensities of FBF1 on the centrosomes in control and LRRC45 depleted cells. Number of centrosomes quantified is 540, 448, 361, and 370 for each respective siRNA treatment. *** p < 0.001 (unpaired Wilcoxon-Mann-Whitney Rank Sum Test). (E) Model for LRRC45’s position in the hierarchy of distal appendages.

Next, we addressed how LRRC45 contributes to appendage formation. LRRC45 depletion did not impact on the centrosome levels of SCLT1 or Cep164 (Figure S4B-D). In contrast, the levels of FBF1 were significantly decreased (Figure 4D). We observed a significant reduction in centrosome levels of Cep123 when Cep123 was detected with a previously published antibody (Sillibourne et al., 2011) (our unpublished observation). However, Cep123 centrosome levels remained unchanged when Cep123 was visualised using our home-made antibody (Figure S6B). We therefore concluded that LRRC45 does not affect Cep123 levels at the mother centriole, yet it may change the way Cep123 interacts with centrioles. Together, our data is consistent with the model that LRRC45 is positioned between Cep83 and FBF1 in the organisation scheme of distal appendage components (Figure 4E).

We next employed the yeast two-hybrid system to determine the interaction network of distal appendage proteins, including LRRC45. For this, we used a combination of full length and truncated constructs to identify their minimal interaction domains (Figure 5B). We reasoned that interactions obtained by the yeast two-hybrid system would reflect direct interactions, as genes encoding for appendage proteins are not present in the yeast genome. Full length LRRC45 interacted with Cep83 and SCLT1 constructs (Figure 5B, red squares). LRRC45 failed to interact with Cep123 or weakly interacted with Cep164 (Figure 5B). LRRC45 C-terminal truncations interacted with truncated forms of FBF1 carrying coiled-coil domains (Figure 5B). Interestingly, the leucine-rich repeats of LRRC45 (LRRC45 1-222 aa) strongly associated with the coiled coil domain of SCLT1 (SCLT1 C-terminus, 554-688 aa) but not with the coiled coil domains of other proteins (Figure S5), indicating interaction specificity to SCLT1. These results, together with the complete yeast two-hybrid data of appendage components (Figure S5), led us to propose that the C-terminal domain of LRRC45 associates with Cep83 and FBF1, whereas the LRRC45 N-terminus binds to SCLT1 (Figure 5C). Cep83 may interact with itself via its C-terminal region (Figure S5 and 5C).

**Figure 5.**
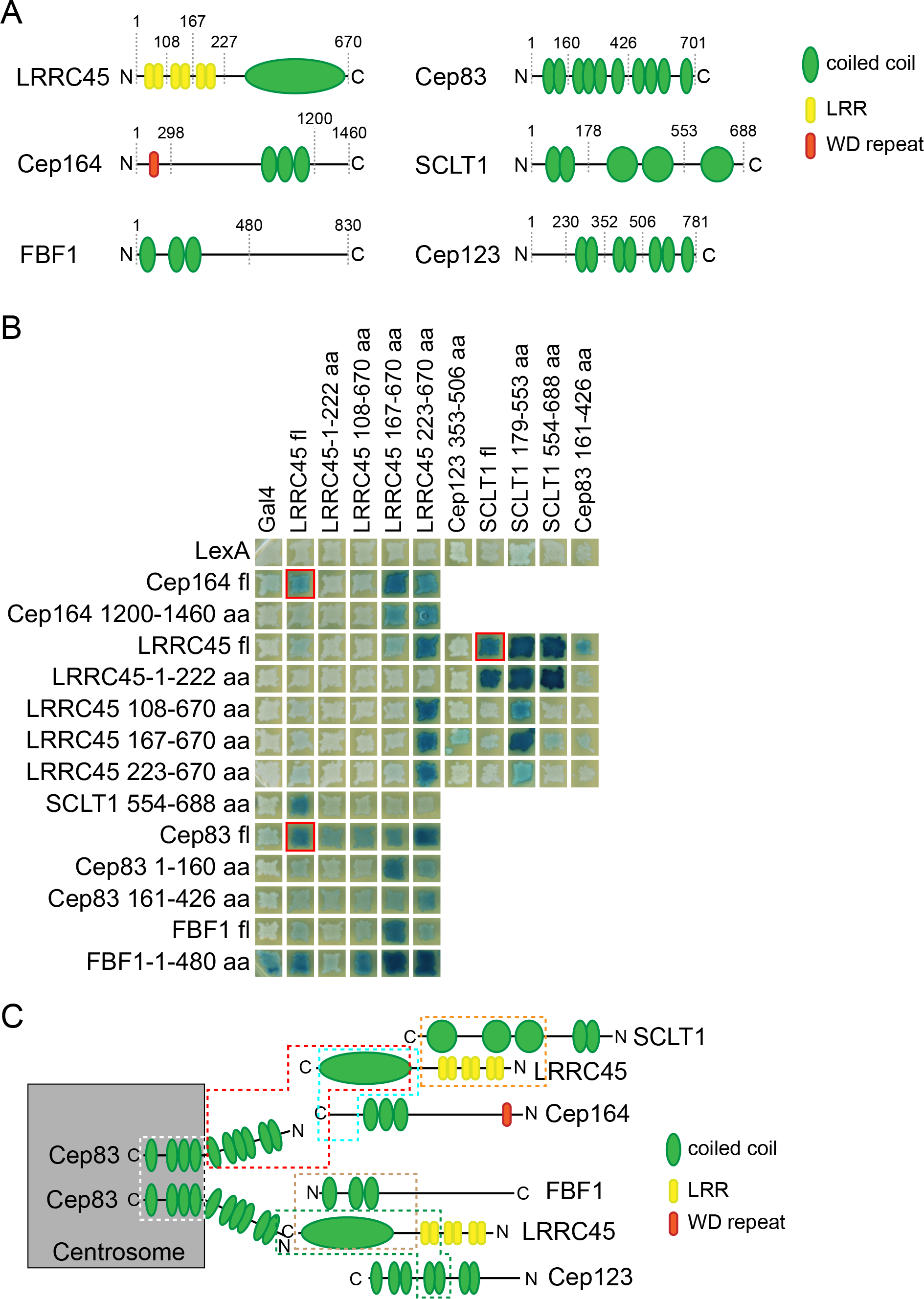
LRRC45 interacts with Cep83, SCLT1 and Cep164 in the yeast two-hybrid system. (A) Schematic representation of existing domains in appendage proteins. Numbers represent amino acid positions. The dashed lines indicate two-hybrid truncated constructs created. (B) Yeast 2 hybrid assay for mapping the interactions between LexA and Gal4 fusion proteins, as indicated. Development of blue colour indicates interaction. Full set of interactions is shown in Figure S5. (C) Model of the interaction map of distal appendages according to yeast 2 hybrid assay. Dashed lines of the same colour encircle interacting domains.

### LRRC45 is required for Rab8-centrosomal localisation but not for CP110 removal

To determine which step of cilia biogenesis is regulated by LRRC45, we first analysed the early ciliogenesis markers EHD1 and Rab8. EHD1 was shown to localise to pre-ciliary endosomal vesicles that accumulate around centrosomes during early steps of ciliogenesis (Lu et al., 2015). The enrichment of EHD1 at centrosomes is independent of Rab8 and vice-versa (Lu et al., 2015). Using a cell line stably expressing mNeonGreen-EHD1, we determined EHD1 protein localisation in control, LRRC45, Cep164 and Cep123 depleted cells stained for Arl13b (cilia marker) and γ-tubulin (basal body marker). Cells were cultured in a serum-free medium for a short period of time to enrich for early stages of ciliogenesis. mNeonGreen-EHD1 was detected as dot-like structures in the cytoplasm or associated with centrosomes (Figure 6A). mNeonGreen-EHD1 was at centrosomes in control cells early in ciliogenesis in which Arl13b accumulation was not yet observed (Figure 6A). Cep164 and Cep123 but not LRRC45 depletion strongly reduced the percentage of cells with docked mNeonGreen-EHD1 at centrosomes (Figure 6A). We thus concluded that Cep164 and Cep123 are mainly required for efficient accumulation of early EHD1 endosomal vesicles at centrosomes. LRRC45 has only a minor function in this early step of ciliogenesis.

**Figure 6.**
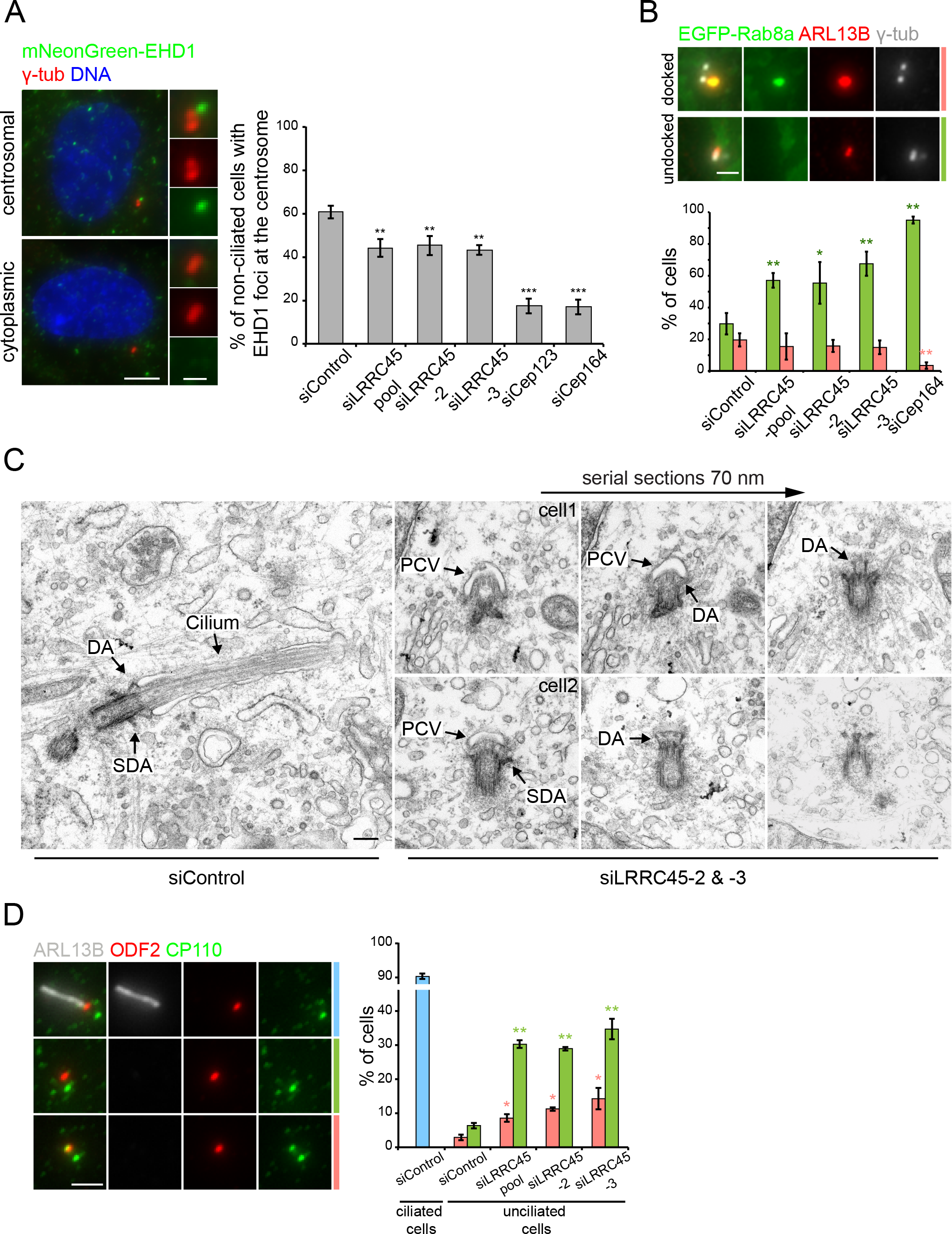
LRRC45 is required for recruitment of Rab8a positive vesicles to the mother centrosome. (A) Representative images of RPE1 mNeongreen-EHD1 cells with centrosomal or cytoplasmic EHD1 (green) foci. Images were taken after 16 h of serum starvation. γ-tubulin (red) and DAPI (blue) serve as markers for centrosomes and nuclei, respectively. Graphs show mean ± standard deviation of mNeongreen-EHD1 positive centrosomes in non-ciliated cells upon respective siRNA treatments (three independent experiments). ** p<0.01 and *** p < 0.001 (Student’s t-test). Scale bar: 5 μm and 1 μm. (B) Representative images of RPE1 EGFP-Rab8a ciliated cells (blue bar) and non-ciliated cells with (red bar) or without (green bar) Rab8a positive vesicles at the centrosomes. Images were taken after 16 h or serum starvation. γ-tubulin (grey) and ARL13B (red) serve as markers for centrosomes and cilia, respectively. The graphs show mean ± standard deviation of the indicated phenotypes after the indicated siRNA treatments. Bars indicate standard deviations of three independent experiments. * p<0.05 and ** p < 0.01 (Student’s t-test). Scale bar 2 μm. (C) Representative electron micrographs showing ciliated WT and non-ciliated but ciliary vesicle docked LRRC45 depleted RPE1 cells (using combination of siLRRC45-2 and siLRRC45-3). DA, distal appendage; SDA, subdistal appendage; PCV, primary ciliary vesicle. Scale bar 200nm. (D) Representative images of RPE1 ciliated cells (blue bar) and non-ciliated cells with (red bar) or without (green bar) CP110 at the mother centriole after 16 h or serum starvation. ODF2 (red) and ARL13B (grey) serve as markers for mother centriole and cilia, respectively. The graphs show mean ± standard deviation of the indicated phenotypes after the indicated siRNA treatments. Bars indicate standard deviations of three independent experiments. * p<0.05 and ** p < 0.01 (Student’s t-test). Scale bar 2 μm.

To analyse Rab8 localisation, we used an EGFP-Rab8 RPE1 stable cell line. In agreement with a previous report (Schmidt et al., 2012), serum starvation led to EGFP-Rab8 accumulation around centrosomes in dependence of Cep164 (Figure 6B). Similarly, EGFP-Rab8 accumulated at centrosomes in control but not in LRRC45-depleted cells (Figure 6B) (Schmidt et al., 2012). This indicates that LRRC45 is required for Rab8-positive vesicle docking at the mother centriole upon induction of cilia biogenesis. Thus, in clear difference to Cep164 and Cep123, LRRC45 is required for Rab8 but not EHD1 recruitment to the basal body in early ciliogenesis.

Together, these results suggest that vesicles dock at the mother centriole without LRRC45. To investigate whether this was indeed the case, we used transmission electron microscopy (TEM) to visualise the basal body of cells depleted for LRRC45. We observed an elongated cilium from the basal body in all control cells (Figure 6D). In the majority of LRRC45 siRNA-depleted cells, the cilium was absent but a vesicle capped the distal end of the mother centriole (Figure 6D). This indicates that small vesicle docking and fusion does not depend upon LRRC45. Rather, LRRC45 is required cilia membrane and axoneme extension.

The CP110/Cep97 complex is a negative regulator of ciliogenesis (Spektor et al., 2007). As Rab8-positive vesicle docking was largely impaired in cells lacking LRRC45, we hypothesised that the subsequent steps of ciliogenesis, including the removal of CP110/Cep97 inhibitory complex from the mother centriole would be inhibited. To analyse whether LRRC45 is required for CP110/Cep97 removal at the mother centriole, we followed CP110 centrosomal localisation upon serum starvation in control and LRRC45-siRNA depleted cells (Figure 6C). As expected, CP110 was absent from the basal body of ciliated control-depleted cells (Figure 6C). In a small percentage of control cells, CP110 removal from the mother centriole (labelled by ODF2) was observed even before the Arl13b-marked ciliary membrane elongated (Figure 6C, siControl green bar). Strikingly, CP110 could not be detected at the mother centriole in approximately 75% of non-ciliated cells lacking LRRC45 (Figure 6C, green bars). The removal of the CP110/Cep97 complex was shown to require distal components including Cep164 and the kinase TTBK2 (Goetz et al., 2012; Schmidt et al., 2012). Consistently, Cep164 (Figure S6D) and TTBK2 centrosomal levels were not affected by LRRC45 depletion (Figure S6F). Our data thus indicate that LRRC45 is dispensable for CP110/Cep97 complex regulation at the mother centriole at early steps of ciliogenesis.

### LRRC45 promotes centriolar satellite localisation

To better understand how LRRC45 contributes to ciliogenesis, we asked whether LRRC45 influenced the microtubule-dependent trafficking of centriolar satellites, which are large protein assemblies required for cilia formation (Craige et al., 2010; Garcia-Gonzalo et al., 2011; Hori et al., 2014; Hori and Toda, 2017; Jin et al., 2010; Klinger et al., 2013; Kurtulmus et al., 2016). For this, we analysed the centriolar satellite proteins PCM1 and SSX2IP. In comparison to control cells, the levels of PCM1 and SSX2IP around centrosomes significantly decreased in the absence of LRRC45 (Figure 7A). Moreover, a similar phenotype of satellite disorganisation was observed when Cep83, SCLT1 and FBF1 were siRNA-depleted (Figure 7B). This data thus suggest that distal appendages have a function in centriolar satellite organisation at the mother centriole.

**Figure 7.**
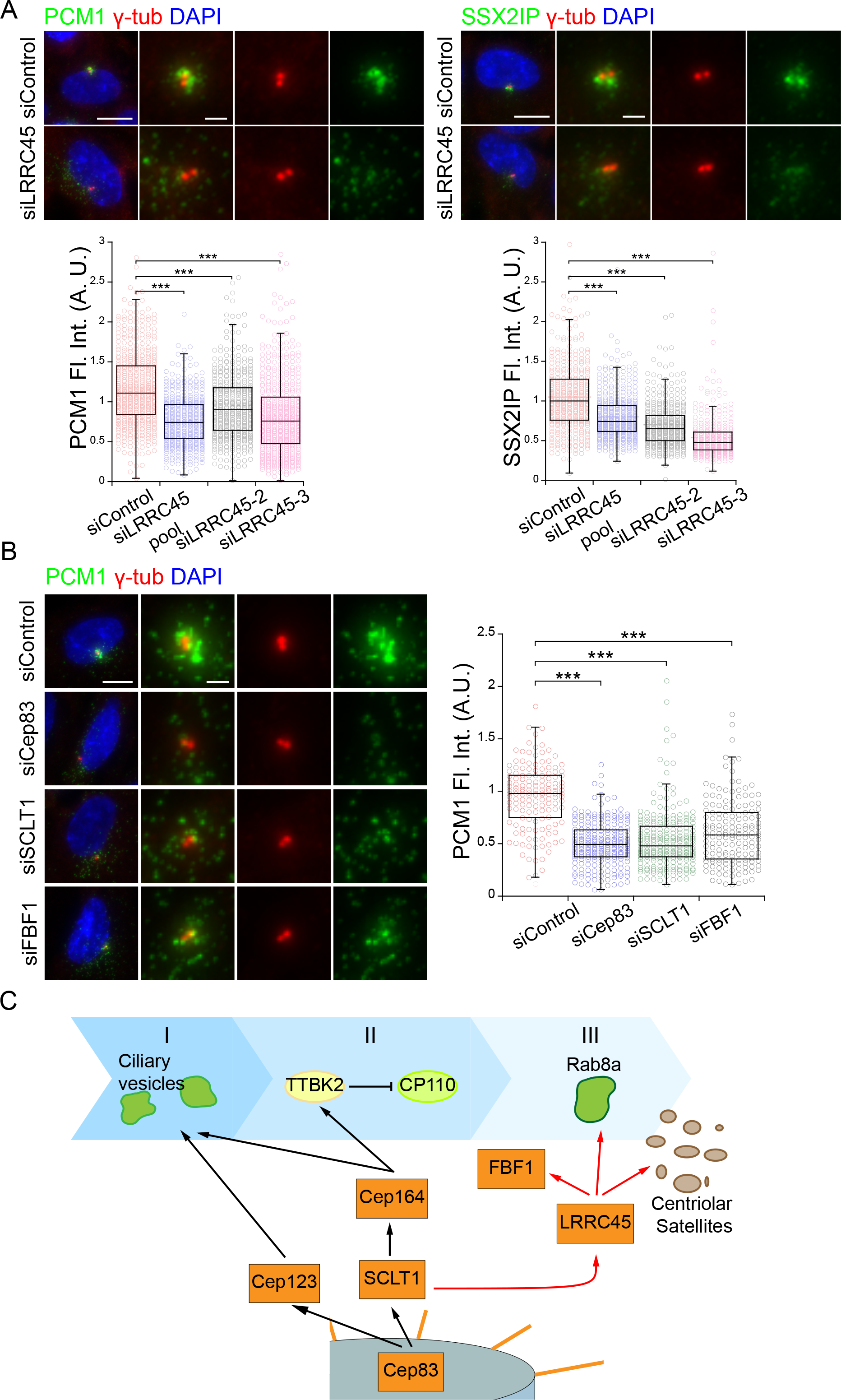
LRRC45 is required for satellite organisation. (A) Representative images show satellite organisation using antibodies against centriolar satellite markers PCM1 and SSX2IP in control- and LRRC45-siRNA depleted RPE1 cells. γ-tubulin (red) and DAPI (blue) serve as markers for centrosomes and nuclei, respectively. The box/dot plots show the relative fluorescent intensities of PCM1 and SSX2IP quantified around the centrosomes from three independent experiments. *** p < 0.001 (unpaired Wilcoxon-Mann-Whitney Rank Sum Test). Scale bar: 10 μm and 2μm. (B) Representative images show satellite organisation using antibodies against centriolar satellite (PCM1) in control-, Cep83-, SCLT1- and FBF1-siRNA depleted RPE1 cells. γ-tubulin (red) and DAPI (blue) serve as markers for centrosomes and nuclei, respectively. The box/dot plots show the relative fluorescent intensity of PCM1 quantified around the centrosomes. *** p < 0.001 (unpaired Wilcoxon-Mann-Whitney Rank Sum Test). Scale bar: 10 μm and 2μm. (C) Model for LRRC45’s role in ciliogenesis. LRRC45 is recruited to the centrosomes via Cep83-SCLT1, and required for proper FBF1 localisation, docking of Rab8a positive vesicles and organisation of centriolar satellites.

To test whether loss of satellite organisation could be a consequence of microtubule loss at centrosomes in response to LRRC45 depletion, we compared the kinetics of microtubule nucleation at centrosomes in control and LRRC45 depleted cells (Figure S7). To visualise microtubules, we followed the microtubule plus-end binding protein EB1 (Mimori-Kiyosue et al., 2000). Cold treatment depolymerised microtubules and decreased centrosomal EB1 (Figure S7). After raising the temperature to 30°C, microtubules re-formed from centrosomes with similar kinetics in wild type and LRRC45 depleted cells (Figure S7). Similar results were obtained for cells lacking Cep83 (Figure S7). This indicates that distal appendage components, including LRRC45, are not required for microtubule nucleation at centrosomes. Furthermore, the centrosomal levels of the intraflagellar transport protein, IFT88, which is transported to the centrosome in a microtubule-dependent manner (Kozminski et al., 1993; Pazour et al., 1998), were not decreased by depletion of LRRC45 (Figure S6G). This indicates that LRRC45 does not impair microtubule-dependent transport to centrosomes in general.

## Discussion

Appendage proteins at the mother centriole play a critical role during the initial steps in cilia formation. Here, we have identified LRRC45 as a component that associates with distal appendages in dependence of Cep83 and SCLT1. Our data suggest that LRRC45 is essential for early phases of ciliogenesis (Figure 7C). Given that LRRC45 localises to the basal body in diverse cell lines, we propose that the function of LRRC45 is conserved from cells that form a single primary cilium, such as stem and differentiated cells, to cells with multiple motile cilia.

Our data are consistent with the localisation of LRRC45 at the mother centriole between the sub-distal appendage component ODF2 and Cep164. Interestingly, using STED super resolution microscopy we were able to resolve Cep123 as two distinct ring-like structures at the mother centriole. This observation is in agreement with a recent report showing that Cep123 forms a layer close to ODF2 and another one closer to FBF1 (Yang et al., 2017). Our data indicate that LRRC45 decorates the space between the two Cep123 rings. Interestingly, we only observed cells with either two Cep123 rings or only one ring more distal than LRRC45. We thus postulate that the Cep123 layer closer to ODF2 might be transient, regulated in a cell cycle dependent manner or modified in such a way that it becomes less accessible to antibody staining. Cep123 and LRRC45 formed rings with a diameter of 240-290 nm, which were significantly smaller than the Cep164 circles. LRRC45 and Cep123 might then be closer to the centriole wall than Cep164. Recently, distal appendage proteins were proposed to form two spatially distinct domains, named distal appendage blades and matrix. According to this model, Cep83, SCLT1, Cep164 and Cep123 form defined conical-shaped blades while FBF1 makes up the matrix by filling the space between the blades (Yang et al., 2017). Depletion of LRRC45 reduced the levels of FBF1 at appendages, indicating that it is a major determinant of FBF1 centriolar localisation. It is therefore tempting to speculate that LRRC45 might be part of the appendage blade complex that is recruited via Cep83 and SCLT1 to help maintain FBF1 at this location.

LRRC45 associates with the proximal ends of mother and daughter centrioles via the linker protein C-Nap1. Proximal and distal pools of LRRC45 co-exist at the mother centrioles and association of LRRC45 with distal appendages does not depend upon C-Nap1. This implies that C-Nap1 and Cep83-SCLT1 do not compete for LRRC45 binding. Recently, the proximity of the daughter to the mother centriole was proposed to be important for ciliogenesis (Loukil et al., 2017). Our data indicate that centriole separation cannot be the underlying mechanism explaining the lack of cilia formation in LRRC45 depleted cells. First, the depletion of LRRC45 did not cause a significant separation of mother and daughter centrioles in RPE1 serum starved cells, even though these cells did not form cilia. Second, in *CEP250* knockout cells, which lose the LRRC45 proximal pool (Figure 1) (He et al., 2013), LRRC45 remained at the distal ends of the mother centriole. In those cells, cilia could be formed in a LRRC45-dependent manner even when centrioles were more than 8 μm apart. Therefore, our data does not show any correlation between cilia loss and distance between centrioles. It rather indicates that LRRC45 plays a key function in ciliogenesis at distal appendages.

Ciliogenesis is a tightly regulated process that occurs at the G1/G0 phase of the cell cycle and involves the conversion of the mother centriole into the basal body. One critical step during this initial phase is the docking of small vesicles at the distal appendages of the mother centriole, which then fuses to generate a large vesicle that caps the distal end of this centriole (Graser et al., 2007; Joo et al., 2013; Kurtulmus et al., 2016; Schmidt et al., 2012; Sillibourne et al., 2013; Sorokin, 1962; Tanos et al., 2013; Ye et al., 2014). Vesicle fusion is promoted by components of recycling endosomes, including the transmembrane tethering protein EHD1 (Lu et al., 2015). Subsequent steps involve membrane elongation, removal of inhibitory components from the mother centriole and targeted delivery of proteins to the basal body (via IFT-and centriolar satellites-dependent transport) to allow axoneme extension (Figure 7C) (Sanchez and Dynlacht, 2016). Interestingly, defects at early steps of membrane establishment block CP110/Cep97 removal and axoneme elongation, implying that these are highly coordinated processes. The common concept is that distal appendages are key components for this coordination, yet how this is achieved on a molecular level remains unclear. Our analyses show that, unlike Cep164 or Cep123, LRRC45 is not required for EHD1 centrosome accumulation or initial steps of ciliary vesicle docking. In the absence of LRRC45, the ciliary vesicle capped the basal body, which is consistent with the fact that EHD1 is involved in vesicle fusion (Lu et al., 2015). However, in cells lacking LRRC45, axoneme and ciliary membrane extension were blocked. The lack of axoneme extension in the absence of LRRC45 was not due to defects in CP110/Cep97 removal, as the majority of non-ciliated LRRC45-depleted cells had CP110 uncapped mother centrioles. This is in agreement with the fact that LRRC45 did not affect the ability of Cep164 to recruit the kinase TTBK2 to promote CP110 centrosomal removal (Goetz et al., 2012). LRRC45 also did not influence the centrosome levels of IFT88, which is required to promote axoneme extension (Sanchez and Dynlacht, 2016).

Our data indicate that the inability of LRRC45-depleted cells to ciliate is mainly related to defective Rab8-dependent ciliary membrane extension and centriolar satellite organisation at centrosomes. In the absence of LRRC45, the levels of the satellite proteins PCM1 and SSX2IP around centrosomes were significantly reduced. Similar results were observed after depletion of Cep83, SCLT1 and FBF1 but not Cep164 (our unpublished observation). Centriolar satellites have been implicated in targeting of Rab8-vesicles to the mother centriole upon induction of ciliogenesis (Hori et al., 2014; Kim et al., 2008; Klinger et al., 2013; Kurtulmus et al., 2016; Lopes et al., 2011; Nachury et al., 2007; Wang et al., 2016). It is thus tempting to speculate that a yet-to-be-identified component required for axoneme extension is targeted to the forming basal body by Rab8-positive vesicles. Alternatively centriolar satellites might be required for centriolar microtubule remodelling to promote axoneme extension after CP110/Cep97 removal. Future research should thus concentrate on understanding how the function of centriolar satellites and Rab8-vesicles are coordinated by distal appendage components at early stages of ciliogenesis.

We propose that appendages can be sub-divided into three categories/modules based on their function in (I) promoting docking of small ciliary vesicles, (II) recruiting TTBK2 and removing inhibitory CP110/Cep97 complexes and (III) promoting Rab8-positive vesicle docking to initiate membrane extension and axoneme elongation in addition to satellite organisation (Figure 7C). Both, the Cep83-Cep123 and Cep83-SCLT1-Cep164 branch of distal appendages, are required for early steps of small vesicle docking (Graser et al., 2007; Joo et al., 2013; Schmidt et al., 2012; Sillibourne et al., 2013; Tanos et al., 2013). Cep83-Cep123 has an additional function in centriolar satellite organisation (Sillibourne et al., 2013), which is not shared by the Cep83-SCLT1-Cep164 complex (Cajanek and Nigg, 2014; Graser et al., 2007; Oda et al., 2014; Schmidt et al., 2012). Without both modules, subsequent steps related to TTBK2 recruitment, CP110 removal, Rab8-vesicle docking and axoneme extension are blocked. In contrast, the Cep83-SCLT1-LRRC45-FBF1 module works downstream in this cascade of events to promote Rab8-vesicle docking as well as satellite organisation (Figure 7C). Cep164 might also assist in this process, given that Cep164 interacts with components of the Rab8-machinery (Schmidt et al., 2012) (Burke et al., 2014). Importantly, the LRRC45 siRNA phenotype is distinct from other distal appendage protein depletions suggesting that centriole appendages are not simply a scaffold for vesicle recruitment but provide a multi-task platform of cooperating, rearranging proteins with defined functions during ciliogenesis.

## Materials and Methods

### Plasmids and reagents

HsLRRC45 (BC014109), HsCep83 (NM_016122), HsSCLT1 (BC128051), OFD1 (BC096344) were amplified from RPE1, and MmLRRC45 (BC023196) was amplified from NIH 3T3 cDNA libraries, respectively. Briefly, total mRNA from RPE1 or NIH3T3 was isolated using RNeasy Mini Kit (Qiagen) according to manufacturer’s protocol, and used for cDNA synthesis using RevetAid First Strand cDNA Synthesis Kit (Thermo Fischer Scientific). HsFBF1 (BC023549) and HsEHD1 (BC104825) cDNAs were purchased from Dharmacon (clones 4109753 and 8143828, respectively). HsCep123 cDNA was a gift from M. Bornens (Sillibourne et al., 2013). Coiled coil domains of the proteins were determined using COILS prediction programme (Lupas et al., 1991), and truncations were cloned into yeast two-hybrid vectors. For rescue of experiments, MmLRRC45 (from mouse cDNA), IRES2 (from pQCXIP-GFP) and GFP (from pEGFP-C1) and cloned into pRetrox-TRE3G (Takara Bio) using NEB HiFi assembly (New England Biolabs). HsEHD1 was cloned into pMSCV-Zeo-C1-mNeonGreen (pMSCV-Zeocin was a gift from David Mu (Addgene plasmid # 75088) (Kendall et al., 2007). More detailed information can be found in Table 1.

### Cell culture and transfection

h-TERT-immortalised Retinal Pigment Epithelial (RPE1, ATCC, CRL-4000, USA) cells were grown in DMEM/F12 (Sigma Aldrich) supplemented with 10 % fetal bovine serum (FBS, Biochrom), 2 mM L-glutamine (Thermo Fischer Scientific) and 0.348 % sodium bicarbonate (Sigma Aldrich). Medium of RPE1 cells stably expressing GFP-Rab8a was supplemented with G418 (500 μg/ml, Thermo Fischer Scientific) (Schmidt et al., 2012). NIH 3T3 cells (ATCC, CRL-1658) were grown in DMEM High Glucose (Sigma Aldrich) supplemented with 10% newborn calf serum (NCS, PAN-Biotech). HEK293T cells (ATCC CRL-3216) and GP2-293 (Takara Bio) were cultured in DMEM High Glucose supplemented with 10% FBS. Adult neuronal stem cells were isolated from sub-ventricular zone of 8-weeks old C57BL/6 mouse and cultured in DMEM/F12 supplemented with 0.236 sodium bicarbonate, % 6 % Glucose (Sigma Aldrich), 50 mM HEPES (Sigma Aldrich), 3.14 μg/ml Progesterone (Sigma Aldrich), 4.8 μg/ml Putrescine dihydrochloride (Sigma Aldrich), 0.37 U/ml Heparin (Sigma Aldrich), 2 % B-27 supplement (Thermo Fischer Scientific), 0.1 % Insulin-Transferrin-Sodium Selenite Supplement (ITSS, Sigma Aldrich), 10 ng/ml fibroblast growth factor-basic (FGF-2, Sigma Aldrich), 20 ng/ml epidermal growth factor (EGF, Sigma Aldrich). All cell lines were grown at 37°C with 5% CO_2_. Transient transfection in HEK293T and GP2-293 cells was performed using Polyethyleneimine (PEI 25000, Polysciences). RPE1 cells were transiently transfected with plasmid DNA by either electroporation using the Neon^®^ Transfection System or TurboFect (Thermo Fischer Scientific) according to the manufacturer’s protocol. Stable cell lines expressing MmLRRC45, and HsEHD1 were generated using retroviral-mediated transduction. GP2-293 cells were transiently transfected with pRetrox-TRE3G-MmLRRC45-IRES2-GFP or pMSCV-Zeo-C1-mNeonGreen-HsEHD1. 48 hours after transfection, supernatant was used to transduce RPE1 cells. Expression of TET-ON-inducible constructs was induced by the addition of doxycycline (Sigma-Aldrich) at a concentration of 1-10 ng/ml. RPE1 CNAP-1 (CEP250) knockout cell lines were described previously (Panic et al., 2015). RPE1 and NIH 3T3 cells were incubated in serum-free medium for 16-48 h to induce cilia formation. At least 100 cells were counted for the quantification of ciliated cells for each experimental condition.

### Antibodies

Antibodies against human LRRC45 were produced against C terminal recombinant protein (240-670 aa) in rabbits and guinea pigs. In brief, 6His tagged antigen was purified from BL21 and used to immunize animals. For purification of the antibodies, GST-LRRC45-C (240-670 aa) was used.

Antibodies against human Cep123 and OFD1 were produced against N terminus of Cep123 (1-230 aa) (Sillibourne et al., 2011) and C terminus of OFD1 (145-1012) (Lopes et al., 2011) in guinea pigs. In brief, 6His tagged antigens were purified from BL21 under denaturing conditions and used to immunize animals. GST-Cep123-N (1-230 aa) and MBP-OFD1-C (145-1012) recombinant proteins were used for purification of the antibodies.

Primary antibodies used for indirect immunofluorescence were rabbit and guinea pig polyclonal anti-Cep164 (1:500 and 1:2000, respectively (Schmidt et al., 2012)); rabbit and g. pig polyclonal anti-ODF2 (1:00 and 1:500 respectively,(Kuhns et al., 2013)); g. pig polyclonal anti-IFT88 (1:250) (Gazea et al., 2016). Mouse monoclonal anti-polyglutamylated tubulin clone GT335 (1:2000) was a gift of C. Janke (Institute Curie, France) (Wolff et al., 1992). Mouse monoclonal anti-Chibby (1:100) was a gift of R. Kuriyama (Department of Genetics, Cell Biology and Development at the University of Minnesota, USA) (Burke et al., 2014). Rabbit polyclonal antibodies anti-PCM1 (1:2000) and anti-SSX2IP (1:250) were gift of O. Gruss (Institute of Genetics, University of Bonn) (Barenz et al., 2013).

Antibodies from commercial sources used in this study are as following; rabbit polyclonal and mouse monoclonal anti-ARL13B (1:500, (Proteintech, #17711-1-AP) and 1:50 (UC Davis/NIH NeuroMab Facility, clone N295B/66, # 73-287, respectively); mouse monoclonal anti-γ-tubulin clone GTU-88 (1:500, Sigma Aldrich, #T6557); rabbit polyclonal anti-TTBK2 (1:1000, Sigma Aldrich, #HPA018113); rabbit polyclonal anti-Cep83 (CCDC41) (1:400, Sigma Aldrich, #HPA038161); rabbit polyclonal anti-SCLT1 (1:250, Sigma Aldrich, #HPA036561); rabbit polyclonal ati-FBF1 (1:500, Sigma Aldrich, #HPA023677); rabbit polyclonal anti-EHD1 (1:100, Abcam, #ab109311); rabbit polyclonal anti-MAPRE1 (EB1) (1:300, Abcam, #ab53358); rabbit polyclonal anti-CP110 (1:300, Bethyl Laboratories, #A301-343A).

### RNA interference

Transfections of siRNA were performed using Lipofectamine RNAiMAX transfection reagent (Thermo Fischer Scientific). For LRRC45 depletion, cells were transfected again 48 hours after the initial transfection. Cycling cells were analysed 48h hours after the second knockdown. For cilia analysis, 16 hours after second transfection, cells were serum starved for 48 hours. Phenotypes observed by LRRC45 depletion were considered to be significant only when it is observed with all siRNAs. Detailed information about the siRNAs can be found in the Table 2.

### Rescue experiments

RPE1 TRE3G-MmLRRC45-IRES2-GFP cells were incubated 6h in 0, 1, 5, and 10 ng/ml doxycycline containing media prior LRRC45 double depletion. Cells were all the time maintained in doxycycline containing media.

### Indirect immunofluorescence and microscopy

Cells were grown on coverslips (No. 1.5, Thermo Fischer Scientific) and fixed in ice-cold methanol at -20°C for 10 min. Cells expressing fluorescent protein fusions were fixed with 3 % PFA for 3 min prior to methanol fixation. Coverslips were coated with 0.1mg/ml Collagen A (Biochrom) for culturing NIH 3T3 cells. For adult NSCs, coverslips were coated with 0.1 mg/ml poly-L-lysine (Sigma Aldrich) and 0.05 mg/ml Laminin (Sigma Aldrich) sequentially. Cells were blocked with blocking solution containing 3% IgG free BSA (Jackson ImmunoResearch), 0.1% Triton X100 (Sigma Aldrich) in PBS for 30 minutes and incubated with primary antibodies at 37 °C for 1h. The cells were then incubated with Alexa Fluor-488, -546, -594, or -647 conjugated secondary antibodies (Thermo Fisher Scientific) together with DAPI (4′,6-diamidino-2-phenylindole) for 45 min at room temperature. All antibodies were diluted in blocking solution. Coverslips were mounted with Mowiol (EMD Millipore).

Images were acquired as Z stacks using either Zeiss Axiophot equipped with 63x NA 1.4 Plan-Fluor oil immersion objective, and Cascade:1K EMCCD camera using Meta. Morph software, or using Zeiss Axio Observer Z1 equipped with 63x NA 1.4 Plan-Apochromat oil immersion objective, and AxioCam MRm CCD camera using ZEN software.

3D-SIM images were acquired as Z stacks using a Nikon Ti inverted microscope equipped with 488nm, 561nm and 647nm laser lines, a Nikon Apo TRIF 100x NA 1.49 oil immersion objective and an Andor iXon3 DU-897E single photon detection EMCCD camera. After acquisition, images were reconstructed using NIS Elements program.

STED images were acquired on with STEDYCON STED microscope platform mounted on Zeiss Axio Imager Z2 with a 100X NA 1.46 oil immersion objective.

All images were taken at room temperature. Proteins fused to fluorescent proteins were visualized by the fluorescence signal. Figures were assembled in Adobe Photoshop and Illustrator CS3 (Adobe).

### Preparation and imaging of brain tissue

8 weeks old C57BL/6J mice (Jackson Laboratory) were sacrificed and brains were extracted and fresh-frozen in Tissue-Tek O.C.T. compound (Sakura FineTek). Coronal sections of 12 μm were cut on a Leica cryo-microtome. Sections were permeabilized with PBS containing 0.1 % Triton X-100 prior to fixation with ice-cold methanol -20°C for 30 min, immunostaining performed with overnight primary and secondary antibody incubations at 4 °C. Sections were imaged on Nikon A1R confocal microscope equipped with Nikon N Apo 60x NA 1.4 λs oil immersion objective, using NIS elements software. Mice were obtained from the DKFZ Center for Preclinical Research facility, housed under standard conditions of 12 hours of light and 12 hours of dark daily and fed ad libitum. All procedures were in accordance with the DKFZ guidelines and approved by the “Regierungspräsidium Karlsruhe”.

### Transmission Electron Microscopy

RPE1 cells were grown on coverslips, rinsed with PBS and fixed with a mixture of 2.5% glutaraldehyde/1,6% paraformaldehyde for 30 min at room temperature. The fixative was washed out with 50 mM cacodylate buffer. After post-fixation in 2% OsO_4_ for 1 h at 4°C, cells were incubated in 0.5% aqueous uranyl acetate overnight, rinsed in water and then dehydrated at room temperature using ascending ethanol concentrations. The coverslips were subsequently set on spurr (Serva) filled capsules and polymerized overnight at 60°C for 48 h. Serial ultrathin sections (70 nm) were post stained with uranyl acetate and lead citrate and examined at a JEM-1400 electron microscope (JEOL, Tokyo), operating at 80 kV and equipped with a 4K TemCam F416 (Tietz Video and Image Processing Systems GmBH, Gautig). Image brightness and contrast were adjusted in ImageJ.

### Microtubule re-growth

Cells grown on coverslips were incubated on ice for 2 hours in order to depolymerize microtubules. Cells were fixed in ice-cold methanol after incubated at 30 °C for 0, 5 and 20 sec., respectively.

### Yeast two-hybrid assay

Full length or partial ORF of the indicated genes were cloned into pMM5 (LexA) and pMM6 (Gal4), and assay was performed as described (Geissler et al., 1996). Yeast growth conditions were as described (Sherman, 1991).

### Image Processing and Analysis

Quantification of fluorescence intensity was performed using maximum projection of images using FiJi and CellProfiler (Carpenter et al., 2006; Schindelin et al., 2012; Schneider et al., 2012). Briefly, images were corrected for background signal by applying a top hat filter, and then centrosomes were segmented using γ-tubulin signal. Identified centrosome areas were expanded by 2 pixels and 10 pixels for intensity measurement of centrosomal proteins and centriolar satellites, respectively. Intensity measurement from replicate experiments were normalized and combined. Inter-centrosomal distances were manually using FiJi. Statistical analyses of fluorescence intensity measurements and ciliation assays were performed using Wilcoxon and twotailed Student’s t tests; significance probability values are: *: P<0.05; **: P<0.01, ***: P<0.001 Statistical tests were performed in Excel (Microsoft) and KaleidaGraph (Synergy Software).

## Author Contributions

Conceptualisation B.K. and G.P.; Methodology, B.K., G.K. and G.P.; Investigation, B.K., C.Y., J.S. and A.N.; Writing – Original Draft, B.K. and G.P.; Writing – Review &Editing, B.K., A.N., S.H., G.K. and G.P.; Funding Acquisition, G.P.; Resources, S.H., G.K. and A.M.V.; Supervision, G.P.

## Acknowledgments

We would like to thank to Marko Panic, Carsten Janke, Michel Bornens, Andrew Fry, Ryoko Kuriyama and Oliver Gruss for sharing reagents; Astrid Hofmann for excellent technical support; Rafael Dueñas-Sánchez for help with antibody production; Berati Cerikan for working on the initial stage of the project and cloning SCLT1 and Cep83; the microscopy facility of the DKFZ, and Ulrike Engel from the Heidelberg-University Nikon Imaging Center for providing access to microscopes, help with imaging and image processing. We are grateful to Abberior Instruments GmbH for the allowing access to STEDYCON STED microscope platform. This project was funded by the collaborative research grant of the DFG (SFB873) granted to G.P. (Project A14). Core funding for microscopy was provided by the SFB873, project Z03. B.K. is a member of the Hartmut-Hoffmann Berling International Graduate School of Molecular and Cell Biology of the University of Heidelberg (HBIGS) and funded by the SFB873. G.P. holds a Heisenberg Professorship from the DFG (PE1883/3).

